# Functional high capacity exosome-encapsulating bioinspired hydrogel promotes microvascular bed expansion in diabetic mice

**DOI:** 10.1101/2024.08.23.609402

**Authors:** Bibi S Subhan, Juan F Cortes Troncoso, Priya Katyal, Michael Meleties, Tamara Mestvirishvili, Fernando Arias, Andrew Wang, Kelly Ruggles, Igor Dolgalev, Paolo Mita, Jin Kim Montclare, Piul S Rabbani

## Abstract

Chronic wounds present a significant clinical challenge due to impaired skin microvasculature, particularly in diabetes. The tissue engineering study introduces therapeutic Exo-Q, a unique thermoresponsive polymer hydrogel created with Q protein nanofibers and cultured human bone marrow multipotent stromal cell exosomes with consistent gene signature. Limited, local and topical application of Exo-Q hydrogel is feasible for maximum neovascularization during murine diabetic wound closure in a xenotransplantation model, as well as compatibility and vascular delivery in human skin in a xenograft model. Exo-Q hydrogel treatment significantly reduces diabetic wound closure time within ranges of non-diabetic wounds. This innovative, non-invasive tissue engineered therapeutic option offers a promising approach to addressing the complex pathologies of non-healing wounds.

## Introduction

The experimental therapy of delivering small extracellular vesicles (sEVs) into chronic skin wounds, particularly diabetic ulcers, has gained significant attention. Chronic or non-healing skin wounds remain open over a month, and are stalled at an inflammatory phase instead of a typical multiphasic progression towards closure(Lazarus et al., 1994). Severely compromised skin microvasculature is characteristic of diabetes and is a major deterrent for molecular interactions that are necessary for the spatiotemporal nature of wound healing in mammalian skin (Okonkwo & DiPietro, 2017). Despite a prevalence similar to that of heart disease and a 5-year mortality rate comparable to that with cancer(Armstrong David G. et al., 2017; Sen, 2023), clinically effective therapies for chronic wounds remain elusive. These statistics reinforce the pressing need for effective therapeutic modalities that predictably address the complex pathologies of non-healing wounds, including severely impaired microvasculature. The EV-conferred healing advantage suggests a promising tissue engineering alternative, as shown by large numbers of recent translational science studies worldwide(Freedman et al., 2023; Subhan et al., 2022).

Small EVs or exosomes are plasma membrane-bound secreted nanovesicles 30-150nm in diameter, that carry RNA, proteins and such biologic cargo from a source cell to affect multiple biochemical and cellular processes upon interaction with recipient cells(Caplan & Correa, 2011; Kalluri & LeBleu, 2020). We and others have used human bone marrow multipotent stromal cells (BMSCs) and their exosomes to successfully promote wound closure in animal models with severely compromised healing(Kuhn et al., 2020; Rabbani et al., 2018; Subhan et al., 2021; Tao et al., 2017). MSC exosome-based therapy for diabetic wounds orchestrates tissue-level change, exhibiting better efficacy than single cytokine approaches (Kehl et al., 2019; Mendt et al., 2019). Safe clinical translation of this relatively new tissue engineering avenue requires scientific rigor, beginning from critically verified BMSC cell culture to EV prep characterization.

Ensuring that exosome preps are potent and available at the diabetic wound site requires optimized delivery methods. Studies administering EVs as treatment mostly report punctate bolus under the skin, though retention and distribution following such injections are highly variable (Brennan et al., 2020; Wiklander et al., 2015). Hydrogels offer potential for consistent delivery, as they may absorb and trap EVs, as they do for small molecules in three-dimensional polymer chain networks (Gao et al., 2021; Safari et al., 2022). We previously reported Q proteins which self-assemble into higher order nanofibers that physically entangle to form a thermoresponsive hydrogel, and importantly, without crosslinkers that may interfere with desired delivery. The pliable Q-hydrogel undergoes a gel-to-solution transition upon reaching mammalian temperatures, a demonstration of upper critical solution temperature (UCST) behavior. It would facilitate our goal of delivering exosome biomolecular cargo to the non-uniform wound surface and cellular milieu of the diabetic wound bed and periphery. These features set Q apart from majority of non-protein hydrogels that undergo solution-to-gel transitions instead at mammalian tissue-relevant temperatures (Mandal et al., 2020). Exosomes carried in a Q hydrogel could offer a clinically translatable material that can reduce the pathological features of chronic or delayed wounds.

This study introduces the integration of two nanoscale building blocks, BMSC exosomes and Q protein fibers, to generate a new entity called Exo-Q hydrogel. Exo-Q hydrogel is a unique therapeutic thermoresponsive polymer hydrogel, with exosomes carried by the nanofibers and not in the matrix and is ideal for topical administration on skin. Its gel-to-solution (UCST) behavior makes it adaptive to the irregular topography of a diabetic wound and responsive to the wound environment while retaining therapeutic potency. We build a rigorous framework for replicability and extensive characterization of source human BMSCs and exosome preps. We demonstrate the impact and feasibility of limited, local topical application of Exo-Q hydrogel for maximum neovascularization during murine diabetic wound closure in a xenotransplantation model, as well as compatibility and vascular delivery in human skin in a xenograft model. Inspired by a clinical need for effective therapy, our study offers a novel non-invasive tissue engineered therapeutic option that reduces diabetic wound pathology – in a bedside to bench, back to bedside approach.

## Results

### Small EV/exosome preps are consistent across human donors

To establish consistency of the cell culture conditions, we characterized the cell surface phenotype and trilineage differentiation capacity of bone marrow-derived MSCs cultured based on their plastic adherence. Using flow cytometry, we phenotypically characterized passage 3 cells. BMSCs were CD31^-^ CD45^-^ Cd11b^-^ CD19^-^ HLA-DR^-^ (**Figure 1a**). Greater than 99% of the cells demonstrated expression of the markers CD44, CD73, CD105, CD90 and greater than 78% had expression of CD106. We detected less than 5% of cells with expression of CD36 (**Figure 1a**). We previously found that the multipotency and progenitor status of mouse BMSCs strongly correspond to their capacity to facilitate wound healing in the LepR^db/db^ type 2 diabetes model(Rabbani et al., 2018). To assess the multipotency of these human cells, we cultured passage 3 cells in maintenance media, osteogenic, adipogenic, or chondrogenic media. Unlike when in control media, the human BMSCs were capable of successful differentiation along these mesenchymal lineages in the respective induction media (**Figure 1b**). Our results demonstrate that phenotype and multipotency are comparable among BMSCs from different human donors.

**Figure 1.**
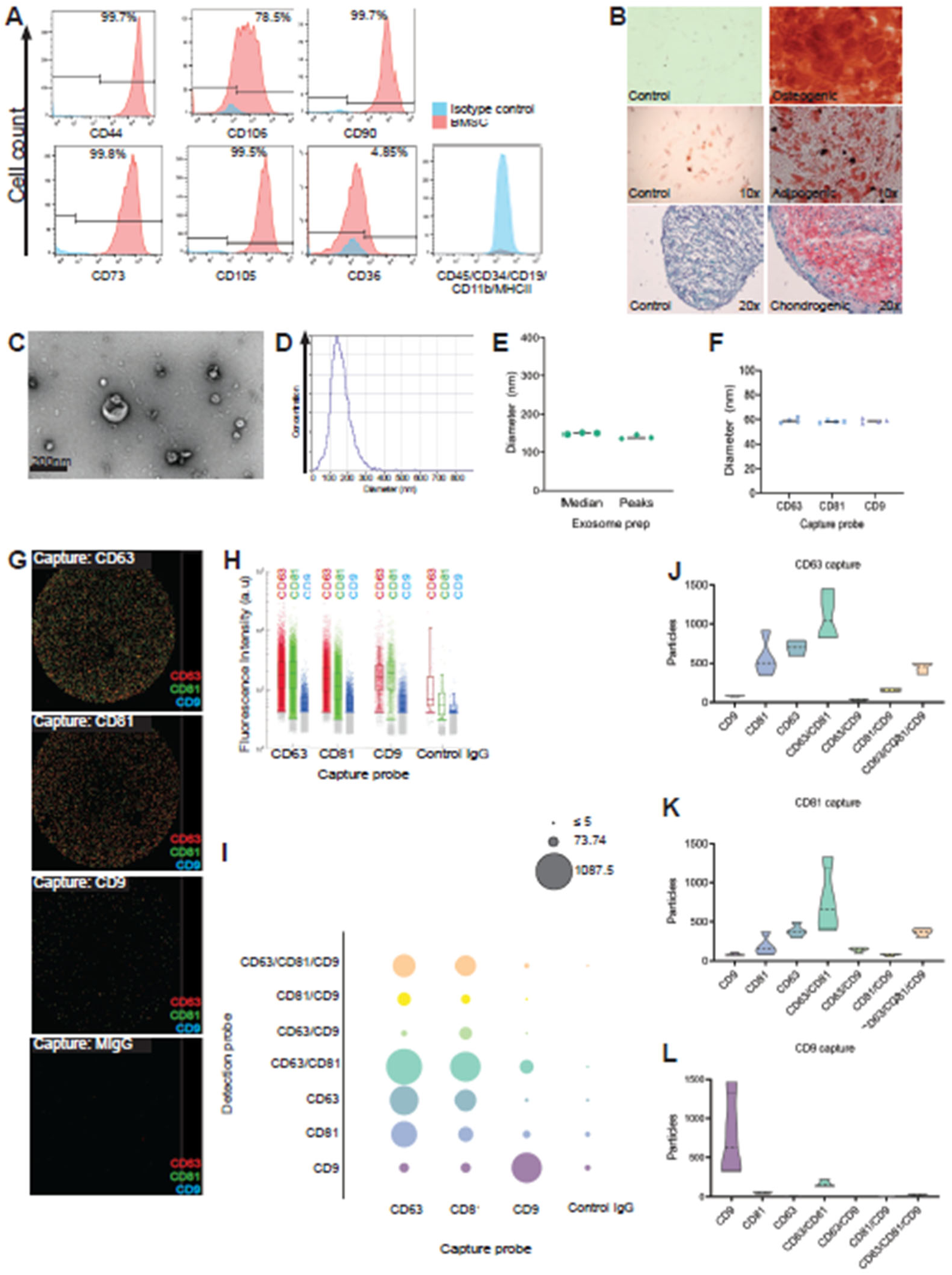
Small EV or exosome preps from cultured bone marrow-derived MSCs are consistent. A) Cell surface marker phenotyping of BMSCs. B) BMSC tri-lineage differentiation capacity analysis. C) TEM of isolated exosome preps. D) Hydrodynamic diameter of exosome preps and quantification (E), using nanoparticle tracking analysis. F) Exosome prep diameter analysis by size interferometry using ExoView. G) Tetraspanin phenotype characterization of exosome preps using ExoView. Each spot has a sandwich of a capture probe, exosome prep, and fluorescent detection probe. Each dot is a nanoparticle. MIgG, control capture probe. H) Abundance of tetraspanins, detected by fluorescence intensity of markers. I) Abundance of colocalization of tetraspanins. J-L) Violin plots distributions of tetraspanin colocalization per capture probe (antibody).

To establish consistency of sEV or exosome preps from BMSC-conditioned media (**Supp. Fig. 1**), we imaged the exosome preps using transmission electron microscopy (TEM) to confirm their presence and size (**Figure 1c**). The TEM preparation also produced the well-documented folding artifact in the exosomes resulting in a collapsed ball or concave shape (Welsh et al., 2024). We then characterized the diameter distribution and concentration using nanoparticle tracking analysis (NTA). Exosome preps from different human donors, but identical cell culture conditions, had similar single modal distributions of EV diameter. Across samples, peaks were between 136nm and 146 nm, while median diameters were between 147nm and 151nm (**Fig 1d-e**). These parameters agree with the guidelines for characterization of sEVs (Welsh et al., 2024). Interferometry sized the exosome preps with average diameters of 59 ± 2.1nm, 58 ± 0.8nm and 58.75 ± 1.7nm for CD63, CD81 and CD9 capture probes, respectively (**Fig 1f**). The exosome preps have near identical diameters with no significant difference across capture probes. Using the Exoview platform, we further phenotyped the exosome preps. Antibodies to the tetraspanins CD63 and CD81 captured maximum number of EVs, while antibody to CD9 captured conspicuously less (**Fig 1g-h**). Colocalization of tetraspanins by detecting captured EVs with fluorescence-labeled antibodies showed prominent populations of CD63^+^CD81^+^ EVs, with minimal CD63^+^/CD81^+^/CD9^+^ EVs or CD63^+^/CD9^+^ or CD81^+^/CD9^+^ EVs in the exosome preps (**Fig 1g, i-k**). Capture with CD9 antibodies demonstrated negligible colocalization with CD63 or CD81 (**Fig 1g, i, l**), suggesting presence of a distinct population of CD9^+^ exosomes. Our results indicate that the exosome preps prepared under the conditions specified contain phenotypically heterogeneous particles from the BMSC conditioned media.

For an additional route of assessment of exosome prep consistency, we performed total transcriptome sequencing of both BMSCs and the exosome preps and established an analysis pipeline (**Fig 2a**). The trimming of the exosome reads is based on the size of small non-coding RNA. The read distribution lengths of the samples show peeks around the expected read lengths of small non-coding RNA(J. Li et al., 2020). We then mapped these trimmed reads to the human genome with the same integrated annotation that is used when generating the counts matrix from featureCounts in MGcount(Hita et al., 2022). In the cells, the reads are not trimmed for the small noncoding RNA, but the mapping to the genome uses the same parameters and integrated annotation as for the exosome samples. Using principal component analysis (PCA) we found that RNA composition varies significantly between source BMSCs and exosome preps with 98% variance (**Figure 2b).** Though we isolated the exosome preps from the very same BMSCs, our results indicate that the packaging of the exosome cargo is specific and reflects a different composition than that in the source cells. The variability among the cells (by donor) or the exosome preps (by donor) is minimal at 1%. To improve the spread of values, we accounted for variable skewness when generating the PCA. Additionally, after regularized log transformation, BMCSCs and their exosomes have a variance of 94%, whereas individual cell populations (by donor) or exosome preps have a variance of 4% (**Supp Fig 2).** A heatmap of expression of significant transcripts shows distinct populations of upregulated and downregulated RNAs in BMSCs and these same RNAs are either upregulated or downregulated in exosomes derived from the very same BMCSs (**Fig 2c**). Notably, the pattern is consistent among the replicates of both the BMSCs and exosomes. The comparison between cells and exosomes reveals a consistent regulation of selective transcription status in the two populations.

**Figure 2.**
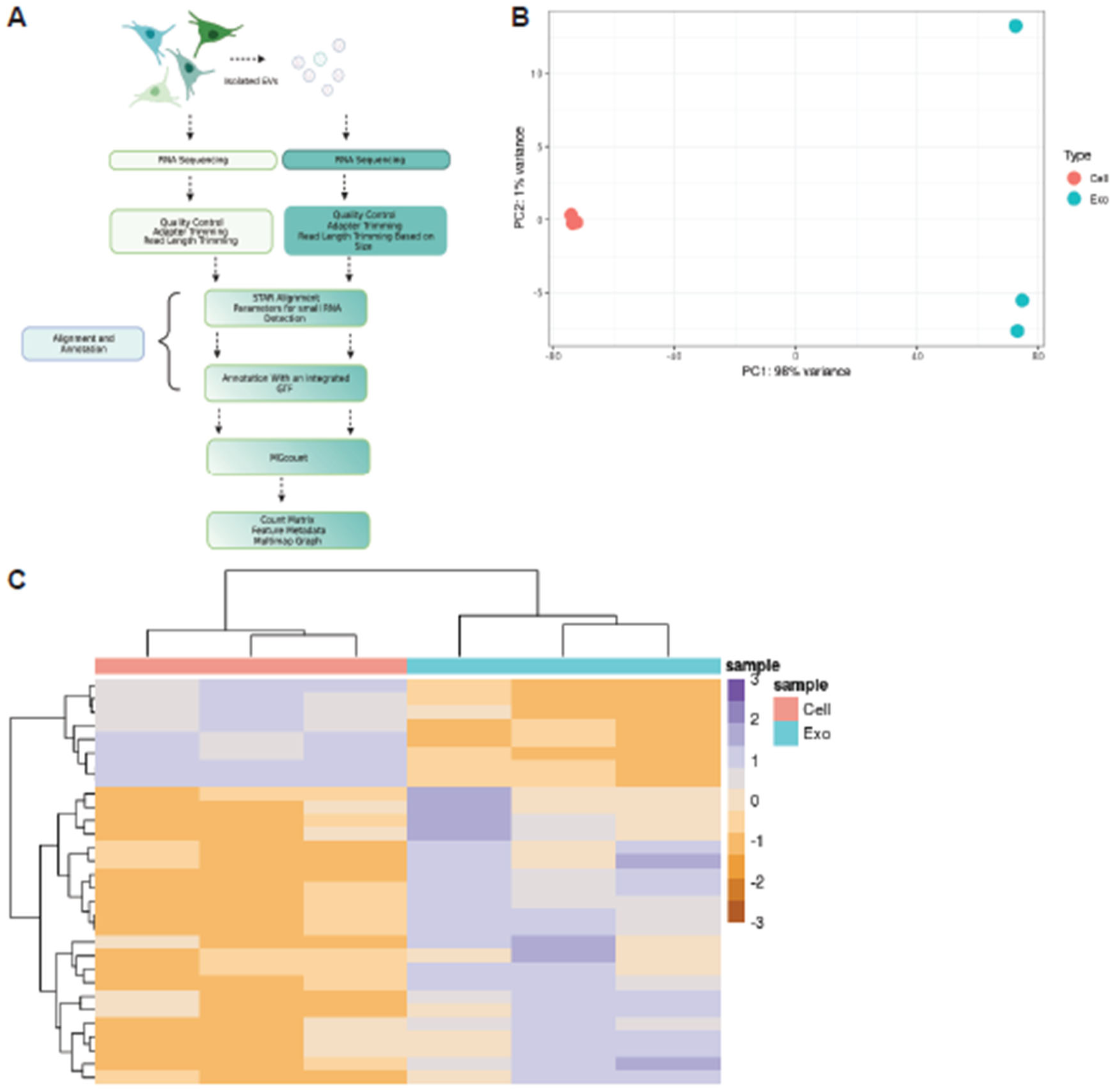
Isolated exosomes have distinct content compared to their source cells. A) Pipeline for RNA seq analysis of cell and exosome samples. B) Variance stabilized PCA plot of cells vs exosomes samples. C) Heatmap of cells vs exosomes of top 30 transcripts with |log2FC| > 0.5 and adj pval <0.1.

We have established that the cell culture conditions and exosome prep isolation methods used are robust and generate replicable exosome preps across BMSCs from different adult human donors. Our exosome preps demonstrate similar diameter distributions, heterogeneous but consistent combination of membrane-associated markers and similar packaging of RNAs from their source BMSCs.

### Exo-Q forms a thermoresponsive flexible hydrogel for maximum contact with wound

For the ideal Q-based exosome delivery platform, we needed a matrix that could encapsulate exosomes in large doses and release them once administered to a mammalian skin wound. Administering exosome preps topically by pipetting, drop by drop, onto the open wounds in LepR^db/db^ type 2 diabetic (T2D) mice did not affect wound closure times, regardless of nanoparticle dosage (**Supplementary Fig 3).** This T2D wound exhibits severe delays in wound healing, when compared to wild type (WT) counterparts and models the chronic nature of human diabetic wounds (Galiano et al., 2004) (**Supp. Figure 3A**, first and second row). We used the validated stented excisional wound model for mouse skin that promotes wound repair by secondary intention and mimics human wound repair and regeneration(Galiano et al., 2004; Rabbani et al., 2018; Short et al., 2023; Subhan et al., 2021).

Q is a self-assembling engineered alpha-helical protein inspired by the coiled-coil domain of cartilage oligomeric matrix protein (COMPcc) Q self-assembles to form nanofibers, which undergo physical entanglement, a form of physical crosslinking, to form hydrogels. We assessed self-assembling properties of Q in the presence of Exo and formulated hydrogels by successfully loading Exo into Q prior to gelation and swelling in water, with minimal impact on the solution-to-gel transition at 4°C (**Figure 3a**). This gave rise to a new hydrogel entity, Exo-Q (**Figure 3b**). To assess topology, TEM image of both Q and Exo-Q hydrogels (diluted to 50 µM) showed that Exo are interspersed within the 3D Q fibrous network, formed of thin and physically entangled nanoscale fibers (**Figure 3c-d**). Nanofibers that we previously reported in Q hydrogel alone(Hume et al., 2014) were also present in Exo-Q hydrogel, confirming that protein self-assembly was not hindered. We then evaluated the mechanical properties or gelation kinetics of Exo-Q using rheology. We observed an increase in storage modulus (G’) (**Figure 3e**) in the presence of Exo. The low tan (delta) for both Q and Exo-Q suggest dominant elastic response. Typically, hydrogels for wound healing applications have transition temperature around 37°C(Lan et al., 2021; Pan et al., 2021). To study the temperature-dependent phase behavior of Exo-Q hydrogels in vivo, we topically applied Exo-Q hydrogels to diabetic wound at post-operative day 1 (POD1). Over the course of 35 minutes, the hydrogel solubilized and went through a gel-solution phase shift at skin wound temperature of 31°C(Gethin et al., 2018) (**Figure 3f**). The hydrogel remained within the confines of the excisional wound margins and did not flow out or over onto the periphery at any time point observed. Our data demonstrates that we have an optimized exosome-loaded hydrogel that can be topically applied to an open wound, has mechanical flexibility and thermoresponsive phase shift that enables maximum proximity of the therapeutic payload to cells of an open diabetic wound bed (**Figure 3g).** The Exo-Q hydrogel matrix enables delivery of exosomes into the wounds without using invasive or painful injection routes, and potentially ensures maximum exposure of exosomes to the wounded skin.

**Figure 3.**
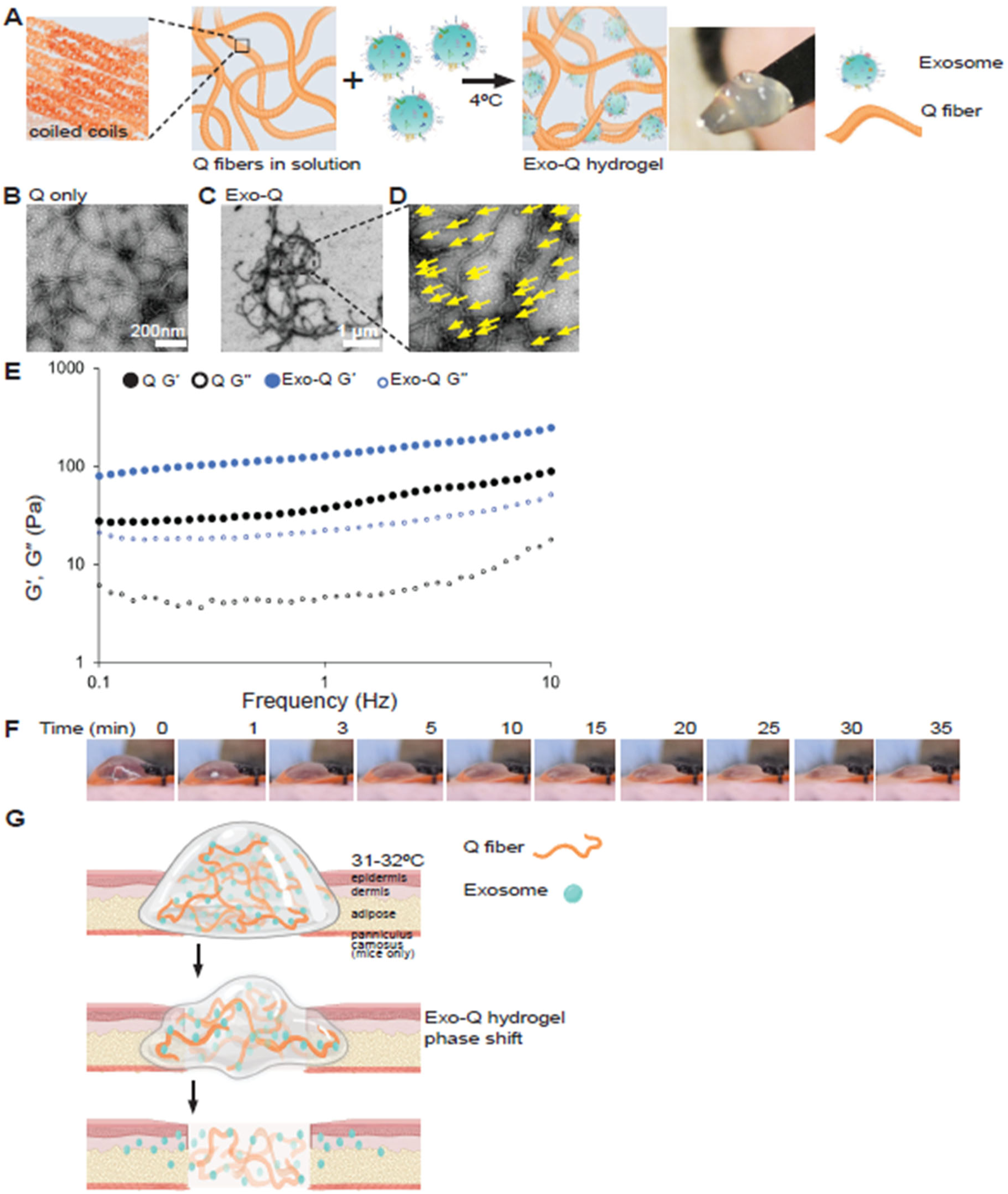
Exosome loaded Q forms elastic thermoresponsive hydrogel. A) Scheme of exosome prep encapsulation in Q fibers to form Exo-Q hydrogel. B-D) TEM of Q only and Exo-Q hydrogels. Yellow arrows indicate exosomes associated with the fibers in inset. E) Rheological properties of Q. Frequency sweeps taken at a strain of 5%. F) Photographs of Exo-Q hydrogel following topical application on diabetic mouse skin wound, over time. G) Schematic representation of Exo-Q hydrogel phase shift and release of exosomes in skin wound.

### Xenotransplantation with Exo-Q hydrogels significantly reduces pathological wound closure in diabetic mice

To analyze pre-clinical therapeutic efficacy of Exo-Q hydrogel, we used a xenotransplantation approach with the Lepr^db/db^ T2D mouse wound model, as in **Supplementary Figure 3**. We administered a single, local, topical dose of Exo-Q hydrogel or control Q-only hydrogel onto diabetic wounds at post-operative day 1 (POD1) and photographically documented the wounds until closure. We observed a dose-dependent effect of exosome prep particles on diabetic wound closure (**Supp. Fig. 4**) and chose to work with 3×10^9^ particles per Exo-Q hydrogel for further experiments. Comparative analysis of the wound planimetric data from the photographs showed that the single Exo-Q hydrogel (3×10^9^ particles) administration resulted in highly significant acceleration of time to closure to 17.67±0.29 days from 25.11±0.99 days for Q-only-treated control and 29.25±0.85 days for untreated diabetic wounds (**Figure 4A-B)**. The median time to closure parallels this pattern; compared to 29.5 days for untreated diabetic wounds and 24 days for Q-only hydrogel-treated diabetic wounds, Exo-Q hydrogel-treated diabetic wounds close in median 17 days, on par with WT wounds (**Figure 4B**). The proximity of the mean and median values for time to closure, for all mouse models and treatment conditions, demonstrates the robustness of the wound model as well as the severe delays in the LepR^db/db^ model (**Figure 4B**). Exo-Q hydrogel-treated diabetic wounds show 83.7% reduction in pathological time to closure compared to that of Q-only treated wounds and 88.9% reduction compared to that of untreated diabetic wounds (**Figure 4B-C**).

**Figure 4.**
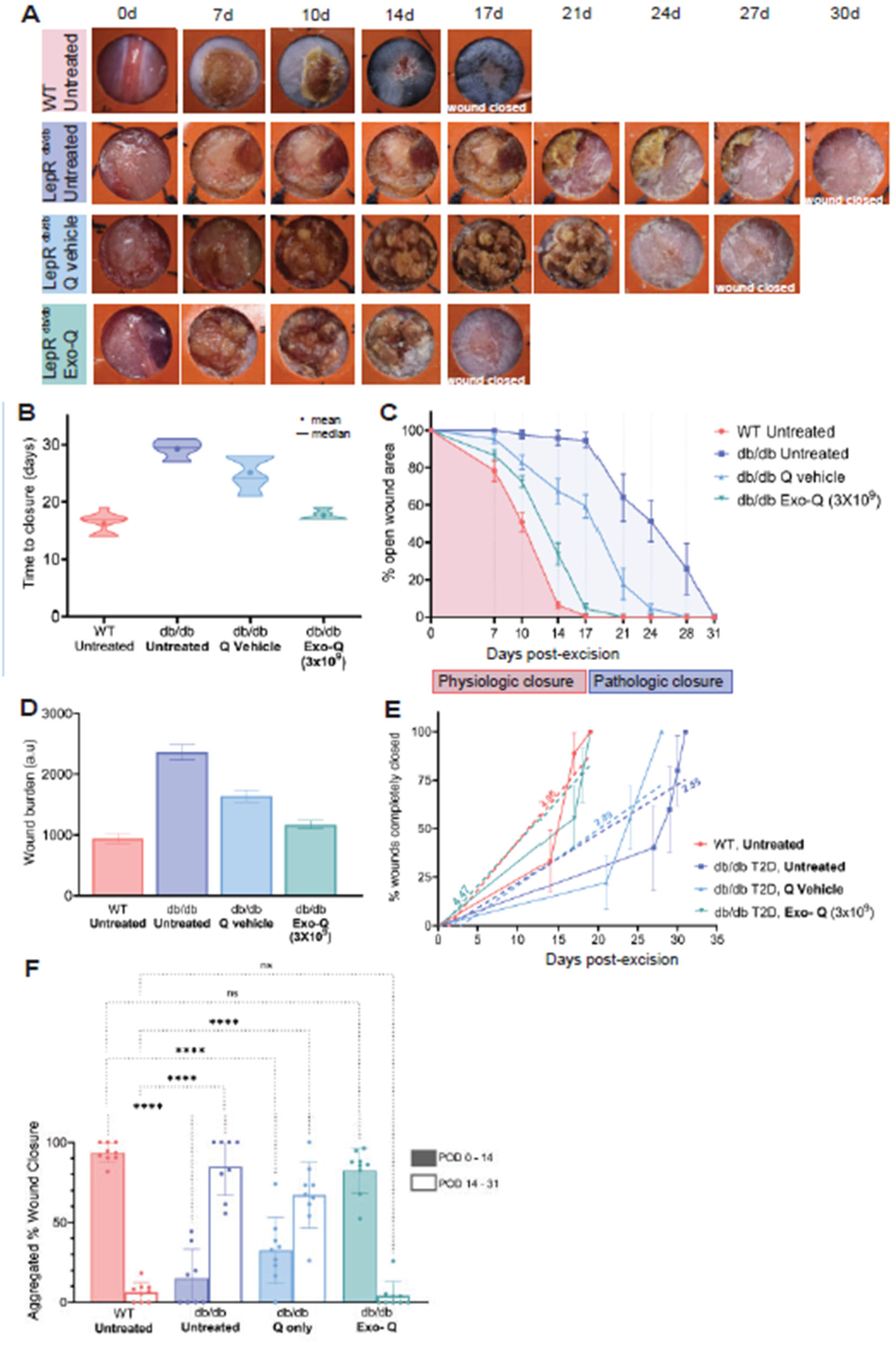
Exo-Q therapy normalizes diabetic wound closure dynamics. A) Time course photographs of WT and LepR^db/db^ excisional wounds, with indicated treatments. B) Wound time to closure statistical analyses. *, p<0.05, ****,p<0.0001. C) Percentage non-healed wound area over time. Red and blue shaded areas, physiologic and pathologic areas under curve, respectively. D) Wound burden, by integrating area under the curve in (C). E) Percentage of wounds closed vs days post-excision. Slope of the dashed regression lines show rate of wound closure. F) Aggregated percentage of healed wound area during POD0-14 and POD14-31, per mouse model and condition. ns, non-significant. ****, p<0.0001.

We then used the quantification of planimetric wound area over time to deduce area under the curve, an indicator of wound closure prognosis(Margolis et al., 2002). The untreated diabetic wounds have the largest area under the curve due to the severe delays in time to closure, compared to the smallest area for WT wounds (**Figure 4C, D**). The area under the curve of the Exo-Q hydrogel-treated diabetic wounds was comparable to that of WT non-diabetic wounds and demonstrates improved healing as well as the reduction in pathological wound burden (**Figure 4D**).

In further comparison of progressive wound closure between treatment groups, WT and Exo-Q hydrogel-treated diabetic wounds have similar wound closure rates (slope of line), while untreated diabetic or Q-only hydrogel-treated diabetic wounds have similar closure rates (**Figure 4E**). This finding is reiterated in analysis of the aggregated percent wound closure at time intervals. The WT and Exo-Q hydrogel groups showed the greater portion of wound closure between POD 0-14, with 93.64% and 82.34% closure, respectively, with no significant difference between these two groups (**Figure 4F**). In sharp contrast, the untreated and Q-only hydrogel-treated diabetic wound groups had only 15.01% and 32.48% closure, respectively, during POD 0-14. For untreated or Q-only-treated diabetic wounds the greater proportion of closure occurred during POD 14-31. Throughout the course of the wound closure studies, blood glucose of LepR^db/db^ mice remained well above 350mg/dL, demonstrating that topical localized application of Exo-Q hydrogels can have efficacy despite persistent hyperglycemia (**Supp. Figure 5**).

To investigate whether wound planimetry data is evident at a histological level, we collected and analyzed wound tissue and immediate peripheral skin at POD5 and POD10 for distance between the migrating epithelial tongues. For best representation of wound epithelial gaps, we only used tissue sections from the wound center axis, dissected in cranial-caudal orientation to align with direction of mouse hair follicle growth. The effect of Exo-Q hydrogel treatment on db/db wound closure is apparent by POD10, but not POD5 (**Figure 5A-B**). At POD5, from the initial 10mm diameter wound, epithelial gaps were 6.06 ± 1.23 mm in WT, 9.34 ± 0.62 mm in db/db untreated, 9.05 ± 1.15 mm in db/db Q-only hydrogel-treated and 8.2 ± 1.48 mm in db/db Exo-Q hydrogel-treated wounds (**Figure 5A-B**). By POD10, the epithelial gaps were 4.95 ± 0.95mm in WT, 8.23 ± 0.68mm in db/db untreated, 8.08 ± 0.71 mm in db/db Q hydrogel-only-treated and 5.60 ± 1.60 mm in db/db Exo-Q hydrogel-treated wounds. The Exo-Q hydrogel-treated db/db wounds demonstrate maximum epithelial regrowth by POD10 in significant difference from untreated and Q-only treated db/db wounds (**Figure 5C**). Our results not only indicate that Exo-Q treatment reverses diabetic wound pathology to a more physiologic healing state, but that xenotransplantation of human BMSC-derived exosomes does not pose as a barrier.

**Figure 5.**
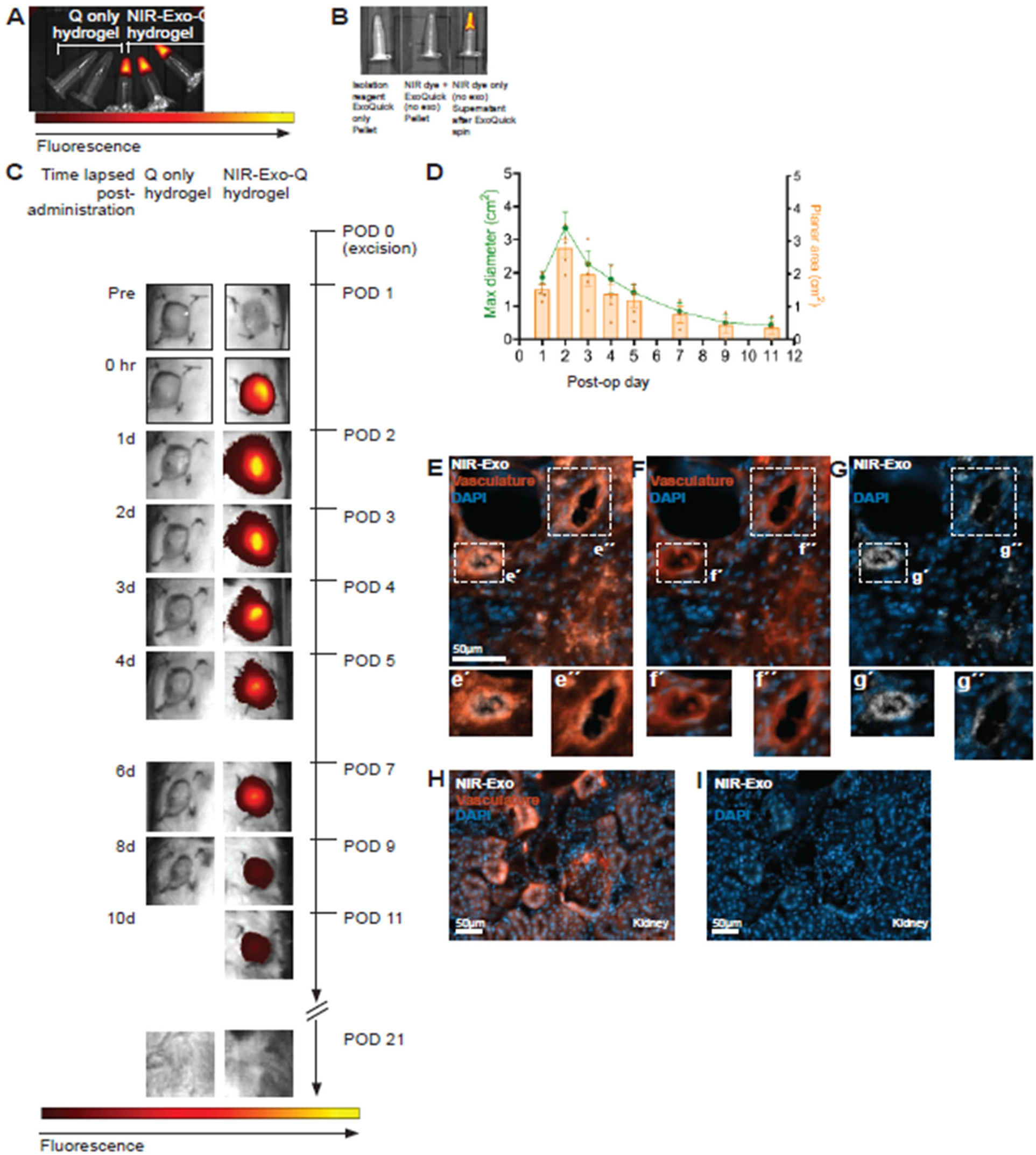
Sustained presence of fluorescence-labeled exosomes in diabetic wounds following Exo-Q hydrogel administration. A) Fluorescence overlaid photograph image of hydrogels, prior to administration. B) Technical controls for fluorescent exosome labeling, demonstrating that unless exosomes are present, the NIR dye does not pellet down with ExoQuick reagent. C) Time course, NIR signal overlaid on photographs of LepR^db/db^ mouse excisional wounds treated with either Q or NIR-Exo-Q hydrogel. D) Quantification of (C). Maximum diameter (green) containing NIR signal and planar traced NIR-area (orange) vs post-operative time, following Exo-Q hydrogel administration. E-G) Wound tissue sections of NIR-Exo-Q hydrogel treated diabetic wound granulation tissue, to detect NIR-Exo. H) Kidney tissue section from same in E-G.

### Exo-Q hydrogel facilitates sustained exosome presence in diabetic wounds

We then assessed in vivo delivery features of Exo-Q hydrogels using NIR fluorescence-labeled exosome preps (**Figure 6A-B**). The NIR label on exosome preps did not hinder Exo-Q hydrogel generation. Following administration of NIR-Exo-Q hydrogels at POD1 on wounds on LepR^db/db^ mice, we used in vivo imaging to longitudinally monitor the NIR epi-fluorescence in the wound bed and immediate periphery (**Figure 6C**). Measurement of the planar area or maximum planar diameter of fluorescent signal (**Supp. Fig. 6**) of the NIR-Exo-Q-treated wounds demonstrated highest readouts 24 hours after administration. Subsequent time points showed diminishing areas of fluorescent signal with no fluorescence detected by POD21 at the latest (**Figure 6C-D**). The fluorescence signal from the wound periphery indicates that the NIR-labeled exosomes spread from the immediate excisional site into the surrounding tissues that contribute to repairing the wound bed. Wound tissue sections from POD7 and lectin-labeled vasculature showed concentration of NIR-Exo signal in endothelial cells, specifically in the cytoplasm (**Figure 6E-G**). We could not detect any NIR fluorescence signals in non-target tissues like the kidneys (**Figure 6H-I**). Our quantitative analysis of fluorescently-labeled exosome presence in Exo-Q hydrogel indicate that the exosomes remain in the targeted skin wound site after topical administration. At the same time, exosomes do not accumulate in non-target internal organs, demonstrating an effective and safe biodistribution profile of Exo-Q hydrogels.

**Figure 6.**
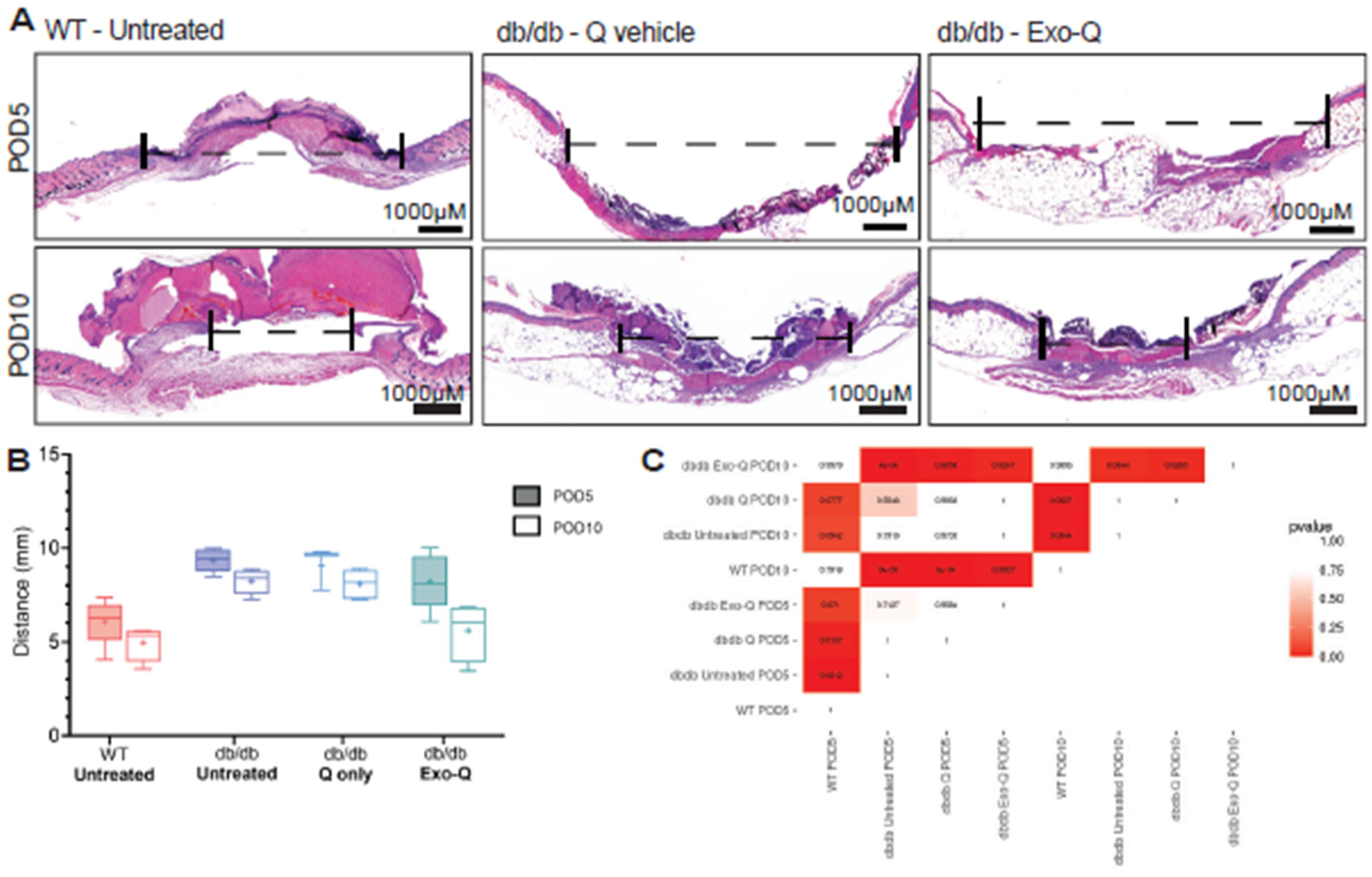
Exo-Q hydrogel administration reduces wound epithelial gap. A) Wound tissue sections from POD5 and POD10 with H&E staining and indicated conditions. Dashed lines indicate gap between wound epithelial edges. B) Quantification and C) significance matrix of epithelial gaps at indicated time points and conditions.

### Wound tissue architecture reflects favorable changes with Exo-Q hydrogel therapy

To analyze the cellular level changes, we used tissue sections and multiplex immunostaining for spatial expression of relevant markers in the wound tissue. Exo-Q hydrogel induced expansion of granulation tissue, a transient vascular-rich connective tissue, in diabetic wounds by POD10, compared to untreated or Q-hydrogel treated ones (**Figure 7A**). This difference is not apparent by POD5 as diabetic wounds regardless of treatment interventions share similar histology (**Supp. Fig. 7**). αSMA expression is highly enriched in the granulation tissue, similar to prior reports(Eckes et al., 2000; Yan et al., 2010). Minimal CD31 and αSMA co-localization demonstrates that majority of αSMA signals in the granulation tissue are not vasculature-associated, but indicate myofibroblasts (**Figure 7B**). Exo-Q hydrogel treatment results in extensive CD31^+^ neovascularization in the αSMA^+^ granulation tissue region of diabetic wounds by POD10 (**Figure 7B**). By POD5, however, all diabetic mouse wounds, regardless of treatment type, lack CD31^+^ expansion (**Supp. Fig. 7A, B**). Expression of αSMA paired with vimentin, a major cytoskeletal protein in dermal fibroblasts and connective tissues(Cheng et al., 2016; Eckes et al., 2000), demonstrates the extent of the transient granulation tissue in both WT and db/db wounds (**Figure 7C**). Co-staining with Ki67, a proliferation marker, revealed proliferating CD31+ cells in the wound peripheral dermis at POD5, as well as Ki67+ vimentin+ cells in both WT and Exo-Q-treated db/db wounds, unlike in untreated and Q-only treated db/db wounds (**Supp. Fig 7 D-E**). Ki67+ Vimentin+ cells were present in the granulation tissue, below the migrating epithelial tongue, in WT mice at POD5. By POD10, colocalization of Ki67 with CD31 or vimentin is rare across WT and db/db wounds, irrespective of treatment type (**Figure 7D-E**). One caveat may be that CD31^+^ endothelial cells are weaving in and out of the tissue and a nucleus may not be visible in the tissue sections analyzed. WT and Exo-Q-treated db/db wounds show similar presence of CD45+ cells in the wound peripheral dermal and adipose layers by POD5, and extensive presence in the granulation tissue by POD10 (**Supp. Fig 7F**, **Figure 7F**). Untreated or Q-only-treated db/db wounds show CD45 accumulation in the wound peripheral skin, but not the granulation tissue (**Fig.7F**). Cleaved caspase 3 (CC3) expression was undetectable at POD5 in WT or any diabetic wound (**Supp. Fig. 7G**), lending support to the safety and lack of adverse responses to the Exo-Q hydrogel. Intriguingly, CC3 is observable in the POD10 granulation tissue in both WT and Exo-Q-treated db/db wounds (**Fig. 7G**). Finally, gene expression of key wound healing-associated factors using whole wound tissue showed significant upregulation of angiogenic and fibroblast-related factors in Exo-Q hydrogel-treated db/db wounds, in comparison to Q-only hydrogel counterparts (**Figure 7H**). Exo-Q-treated db/db wounds showed 1.246 vs 1.002 fold change in *VEGF* expression, 2.214 vs 1.027 fold change in *SDF1* expression, and 1.352 vs 1.005 fold change in *PDGF* expression. This result supports the expansion of granulation tissue evident in tissue section analysis of the Exo-Q hydrogel-treated db/db wounds.

**Figure 7.**
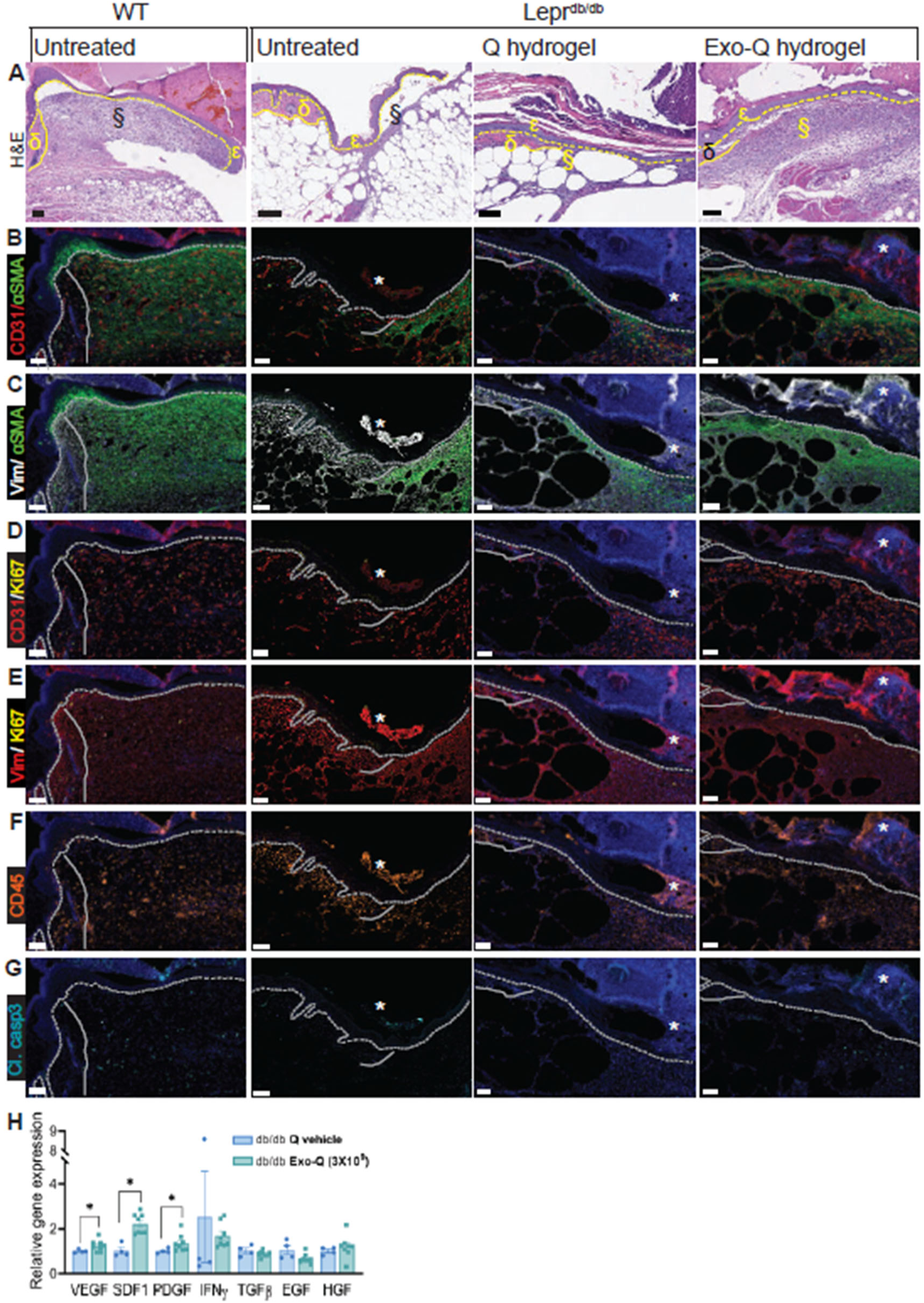
Granulation tissue and wound adjacent dermis are similar between Exo-Q hydrogel-treated diabetic and WT mice. Wound tissue sections from POD10, stained with H&E (A) and multiplexed markers as indicated (B-G). Dashed lines, epidermal-dermal boundary. Solid white line, granulation tissue-dermis boundary. Ɛ, epidermis. Δ, wound-adjacent dermis. §, granulation tissue. *, autofluorescence. Scale bar, 200µm. (H) Relative gene expression of wound healing-associated factors in diabetic wound tissue. *,p<0.05.

### Exo-Q hydrogels demonstrate biocompatibility with human skin

To verify whether Exo-Q hydrogels are compatible with human skin, we used a human skin xenograft wound model. This approach enables studying the human BMSC Exo in Exo-Q hydrogels in allotransplantation (human skin wound), in addition to the xenotransplantation use (mouse skin wounds) thus far. The human skin xenograft contains epidermis and dermis, but with excised adipose tissue, and human hair regrowth in the xenograft indicated vasculature inosculation between human and mouse skin (**Supp. Fig. 8A**). Tissue sections demonstrated the interface between mouse and human skin, as well as the wound in the human skin xenograft (**Figure 8A-B)**. At day 0 of wounding, bleeding following excision further confirmed vascular anastomoses between the human xenograft and mouse skin (**Figure 8C**). Upon topical administration, the Exo-Q hydrogel sank into the wound bed of the thicker human skin (**Figure 8D**), in notable difference from the dome-shaped hydrogel on mouse wounds (**Figure 3F**). The xenograft wound progresses towards wound closure (**Figure 8E**). In vivo imaging of GFP-labeled exosomes in Exo-Q hydrogel detected a fluorescent signal in the human xenograft wound until POD5 (**Figure 8E**). Using a human tissue explant wound model, we applied CD81-GFP labeled exosomes (from human adipose tissue derived MSCs) in Exo-Q hydrogel. Echoing the results in the mouse models, the GFP label signal is likely present in the vasculature of the explant (**Supp Fig. 9**). Our results demonstrate that Exo-Q hydrogel does not pose a barrier to human skin wound healing in an allogenic setting and shows no interspecies discrepancy in biocompatibility.

**Figure 8.**
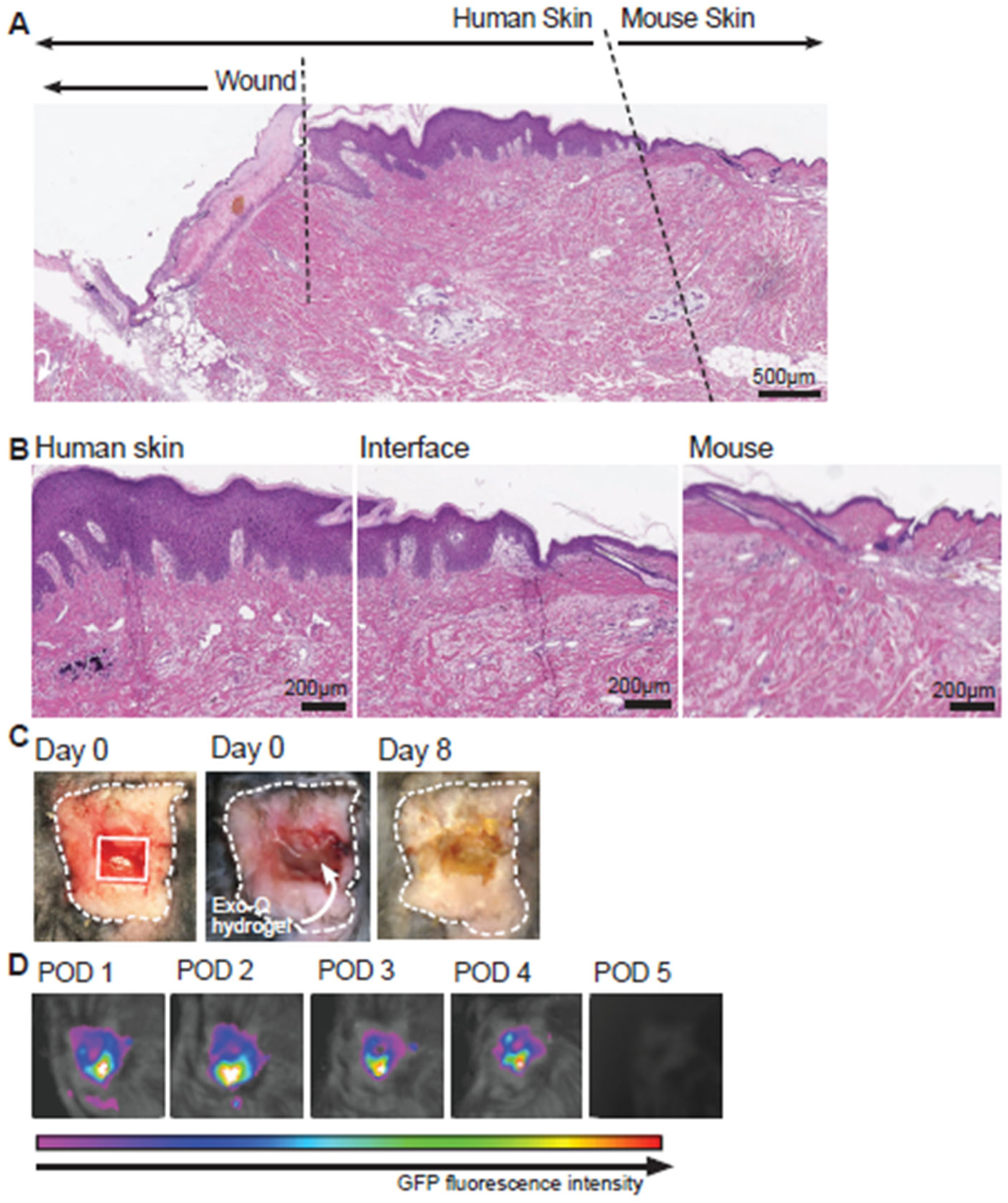
Exo-Q hydrogels are compatible with human skin. A) H&E tissue section of human and mouse skin junction in xenograft mouse model. B) Higher magnification images as indicated. C) Pre- and post-administration of Exo-Q hydrogel into wound in human xenograft. D) In vivo imaging of GFP-labeled Exo in Exo-Q.

## Discussion

Chronic non-healing diabetic ulcers often lead to lower limb amputations, accounting for nearly 70% of non-traumatic amputations as well as 25% of hospitalizations in the US, and are a leading cause of disability worldwide (Armstrong et al., 2023; Zhang et al., 2020). Current standards of care involve the following: assessing disease and social history, repeated invasive measures such as physical or biochemical debridement, weight off-loading, antibiotics and frequent dressings. Permutations and combinations of these approaches lower patient adherence(Osterberg Lars & Blaschke Terrence, 2005; Sen, 2023). While innumerable products and devices, including hydrogels, are available for safe management of chronic wounds very limited numbers are indicated for efficacy of addressing pathology (Verma et al., 2022). Approaches that facilitate easy and limited administration of EV-based therapies have the potential for clinical adaptation and are critically needed to improve the quality of life for patients with diabetes and for predictability of wound treatment outcomes. We developed and optimized a unique and novel topically administered exosome-encapsulating hydrogel, Exo-Q, that can push past the stalled nature of severely delayed diabetic full-thickness skin wounds to accelerate progression of wound healing phases to closure.

Exosomes do not have a singular effect but affect multiple networks of biochemical and cellular processes for significant tissue level impact(Van Delen et al., 2024). Exosomes nullify the risk of unwanted growth and unknown fate of stem cell transplantation, are a major mediator of paracrine or trophic functions of MSCs without inducing adverse immune responses in allogeneic and xenogenic use, and have potential as an efficacious biologic with long shelf life(Caplan & Correa, 2011; Domaszewska-Szostek et al., 2019; Jeong et al., 2011; Sivanantham & Jin, 2022). Meta-analysis of clinical studies to date highlights the safety of BMSC EVs, particularly in allogeneic use(Van Delen et al., 2024). Relatively easy MSC isolation, cell culture and EV prep have contributed to the worldwide increase in tissue engineering studies with EVs(Subhan et al., 2022). In our approach, the human BMSCs or exosomes do not undergo intrusive manipulations. Despite the heterogeneity among small EVs or exosomes, our culture parameters condition the human BMSC and their exosomes for consistent output and minimize batch effects across different donors, as our analysis of RNA content between the source BMSCs and exosomes showed (**Figures 1-2)**. Reporting replicability of MSC-derived exosomes is of paramount importance for clinical translation, as suggested by multiple international bodies(Witwer et al., 2019; Welsh et al., 2024). In this study, we analyzed all RNAs in the BMSCs and EV preps to establish new means of assessing RNA-based signatures of therapeutic EVs.

Despite the skin providing easy anatomic access for wound treatment, the complex histology and diabetic pathology pose challenges for effective delivery and exosome availability. Most tissue engineering studies using EVs, particularly in the context of wound healing, report repeated skin trauma with injections(Brennan et al., 2020; Wiklander et al., 2015). Hence, feasible clinical administration was a critical early factor in our design of an exosome-carrying hydrogel. We have used a self-assembling protein-based hydrogel that gels at lower temperatures and undergoes a gel-to-solution phase shift upon reaching the mammalian skin temperature (UCST). These features are unlike majority of hydrogels in peer-reviewed literature that are non-protein-based, require cross-linking (that can introduce complications) and undergo a solution-to-gel transition at mammalian internal temperatures (LCST) (Q. Li et al., 2021; Shi et al., 2017; M. Wang et al., 2019; Mandal et al., 2020). In fact, numerous pre-clinically studied and the only FDA approved hydrogels are all injectables that gel at the injected site(Chen et al., 2019; K. Wang et al., 2022; Zhou et al., 2019; Mandal et al., 2020).

In contrast to the diffusion-based loading after hydrogel formation in our previous work and the existing literature (Hill et al., 2019; Amengual-Tugores et al., 2023), exosomes and Q fibers in solution together give rise to a new entity, Exo-Q hydrogel (**Figure 3**). We found that Exo-Q hydrogels form using exosomes from both bone marrow and adipose-derived MSCs with equal efficiency (**Suppl. Fig. 9**), indicating potential for expanding the cell sources used for generating exosome preps. In Exo-Q hydrogel, the exosomes are not in the hydrogel matrix but associated with the entangled Q protein fibers, as visualized by TEM (Figure 3). Our visual confirmation of exosome localization within the hydrogel is in exception to majority of reports of exosome-loaded hydrogels, including those in wound healing studies. The incorporation of exosomes, in fact, increases the elasticity of Exo-Q hydrogel compared to Q-only hydrogel. The elasticity imparts ease of topical application of the Exo-Q hydrogel to open skin wounds. Particularly in clinical settings, ease of application may reduce the need for specialized medical training and even be administered by patients themselves to increase adherence to therapy. The elasticity of Exo-Q hydrogel also confers the capacity for the gel to conform to uneven wound beds, including changes in shape due to physical or mechanical changes in the skin. The Exo-Q hydrogel showcases functional exosomes for optimal impact in the wound bed.

A single dose of Exo-Q hydrogel demonstrated highly significant impact on promotion of closure of type 2 diabetic mouse wounds (**Figure 4, 5, 7**), compared to untreated or Q-only hydrogel-treated wounds. We used the LepR^db/db^ mouse to model type 2 diabetes and the severe delays in full-thickness wound closure. This model is widely accepted for translationally relevant pre-clinical studies, as well as its capacity to respond to interventions (Brem & Tomic-Canic, 2007; Galiano et al., 2004; Subhan et al., 2022). We chose to administer Exo-Q hydrogel at POD1 to correspond with the critical acute period after debridement of a chronic wound in the clinic. The Q fibers, while associating with exosomes in Exo-Q hydrogel, do not impede wound healing as evident from scab formation, regeneration of the epidermis and repair of the dermal layers.

Instances where the hydrogel material affects wound closure can confound analysis of the function and effect of exosomes(Yang et al., 2020). Our results suggest that Q fibers in the hydrogel degrade within a time frame suitable for pre-clinical mouse wound healing, as well as human wounds. Importantly, the wound closure effect of the BMSC exosomes delivered in Exo-Q hydrogel-treated diabetic wounds is remarkable and distinct from the lack of differential wound healing phenotype with Q-only hydrogel, as compared to untreated diabetic wounds (**Figure 4**). The efficacy of a single Exo-Q hydrogel dose in this preclinical model is promising for increasing patient adherence with a limited clinical dosage regimen.

For wound healing, the wound bed is not the immediate “gap” but includes the surrounding and recruited cells in the periphery of the gap. We do not observe significant changes at the histological level by POD5, but the Exo-Q hydrogel-treated diabetic wounds begin reflecting the expression patterns in the granulation tissue particularly of WT wounds by POD10 (**Fig. 5**, **Fig. 7 and Supp. Fig. 7**). Though the time to closure data shows that Exo-Q-treated diabetic wounds and WT wounds close in comparable time frames and that majority of closure occurs in the first 14 days (**Fig. 4**), the histology and immunostaining reveal that Exo-Q hydrogel-treated diabetic tissue still has a lag. The decrease in open or non-healed wound area, proxied by scab or non-epithelialized area, is a result of both epithelial growth and contraction from myofibroblasts. Multiplexed immunostaining confirms the presence of expanding regions of a-SMA staining granulation tissue by POD10 in Exo-Q hydrogel-treated diabetic wounds, prognostic of wound closure. While Exo-Q hydrogel treatment of diabetic wounds does not recapture WT wounds at the time points analyzed, it is sufficient to reduce pathological index and have similar healing trajectory (**Figure 4, 5, 7**).

The detection of sustained signal from fluorescently-labeled exosomes in the “gap” and immediate periphery in Exo-Q hydrogel-treated diabetic wounds suggest that the exosomes are present in the targeted anatomical site. However, whether and when exosomes released from the hydrogel disperse further than the fluorescent region remains unknown due to limitations in detection of fluorescence. IVIS or confocal microscopy do not yet have the sensitivity to detect single exosomes with sizes in the order of nanometers. Despite these limitations, we find that Exo-Q hydrogel enables accumulation of BMSC exosomes in the cytoplasm of endothelial cells in the granulation tissue. This outcome is promising, especially in the context of endothelial cells being notoriously difficult to delivery new materials into, even in vitro(Hunt et al., 2010). Confirming our expectations, in sharp contrast to Q-only hydrogel treatment, Exo-Q demonstrated extensive neovascularization in large areas of granulation tissue with corresponding upregulation of angiogenesis associated genes. Our results indicate that treatment of diabetic wounds with Exo-Q hydrogel can promote neovascularization and adequate blood supply to the diabetic tissue. Our results reinforce the argument that administration route of EVs affects their distribution (Wiklander et al., 2015). The premise of chronic wound care for a patient with diabetes includes promoting lifestyle changes, such as glycemic control, as persistent hyperglycemia stymies proliferation(Eming et al., 2014). In our study, despite persistent hyperglycemia as a microenvironmental factor, Exo-Q hydrogel promotes local proliferation of wound bed cells and does not have diminished efficacy.

Though LepR^db/db^ mouse models are validated for wound healing studies and considered evidence for translation, compatibility of a novel therapeutic biomaterial with human skin tissue is not a guarantee. Preclinical administration routes typically demonstrate proof of principle, such as pipetting on a solution rather than feasibility with clinically defined needs. Mouse dorsal skin is 400-700um (Wei et al., 2017) while human skin in common chronic wound areas (feet) is 2-6 mm (Chanda & Singh, 2023). Variation of skin thickness with anatomical location has been a limiting factor for delivery approaches like microneedles (Wei et al., 2017). Following topical administration to full-thickness wounds in xenografts, Exo-Q hydrogel does not hinder wound healing progression in human skin and the fluorescently-labeled exosomes are detectable locally for multiple days (**Figure 8**). The depth of human skin is highly suitable for direct topical administration and spread of the elastic Exo-Q hydrogel into the uneven spaces of the wound upon gel-to-solution phase shift and exposure of the wound bed to exosomes. The Exo-Q hydrogel sinks into the skin “gap” in the thickness of human skin and is conspicuously different from the initial hydrogel “dome” on mouse skin before phase shift (**Figure 3**),

Employing a multidisciplinary approach encompassing in vitro assays, animal models, and translational assays, we provide compelling evidence supporting robust, consistent EV generation across human donors and the efficacy of EV-mediated therapy coupled with advanced delivery biomaterials in chronic wound therapy. Our findings not only underscore the therapeutic potential of this innovative paradigm but also elucidate the underlying mechanisms driving its regenerative effects. We explore the translational implications of our results, laying the groundwork for future clinical trials and interventions. Our approach can be extended to other EV sources, like lipoaspirate MSCs. Cell-free Exo-Q hydrogel have potential for rapid translation as an “off the shelf” option with ease of application and lack of immunogenicity in recipients. Application can extend to instances requiring sustenance or promotion of microvascular beds, such as cardiac repair, or any surgical interventions where sufficient blood supply has to be restored before any further reconstructive options can even be considered. Additionally, Exo-Q hydrogels could have veterinary applications. Future versions will optimize use of chemically defined medium to eliminate FBS, as well as incorporate drugs or antibiotics for multi-targeted therapeutic hydrogels.

The integration of sophisticated delivery biomaterials amplifies the therapeutic efficacy of EV-based therapies. These biomaterials serve as versatile carriers, facilitating the targeted and sustained presence of EVs at the wound site. By providing a conducive microenvironment for EV-mediated regeneration and repair, these biomaterials mitigate barriers to wound healing and foster closure. Their customizable properties will allow for tailored approaches, optimizing therapeutic interventions for broad range of wounds and injuries.

## Materials and Methods

### Cell culture and conditioned media collection

We acquired cryopreserved iliac crest bone marrow-derived multipotent stromal cells from RoosterBio (Frederick, MD). At acquisition, the cells had undergone 6-10 population doublings. We seeded 1×10^6^ cells in a monolayer in 150mm tissue culture plates as passage 1, and incubated in 15% FBS, 1% penicillin-streptomycin, 1% non-essential amino acids in alpha-MEM at a humidified 5% O_2_, 5% CO_2_, 37°C. At passage 3, we washed 70% confluent monolayer of bone marrow-derived multipotent stromal cells with 1xPBS containing Ca2+/Mg2+ to minimize cell signaling disruptions. We added media constituted with EV-free FBS (Gibco, ThermoFisher, MA) and returned the plates to the humidified incubators with the parameters described above. Forty-eight hours after addition, we harvested the conditioned media for processing into the exosome prep.

### Flow Cytometric Analysis of BMSCs

At Passage 3, we used trypsin/EDTA to detach human BMSCs and suspended the cells in 1xPBS/5% FBS for immunostaining. We stained cells for 30 minutes with fluorophore-conjugated antibodies against CD31, CD45, CD11b, CD19, HLA, CD44, CD73, CD90.2, CD105, CD106, and CD36 or isotype (Biolegend, San Diego, CA) and washed 3 times with 1xPBS/5% FBS. Anti-rat, anti-hamster, and anti-mouse, IgK/Negative Control Compensation Particles Sets (Becton Dickinson, NJ) were negative technical controls. We then analyzed immunostained BMSCs on a ZE5 Cell Analyzer (Bio-Rad, CA) and processed with Flow Jo (OR).

| Vendor | Ab and color | Catalog # | Clone |
| --- | --- | --- | --- |
| Biolegend | Anti-CD90 Brilliant Violet 711 | 328139 | 5E10 |
| Biolegend | Anti-CD105 Brilliant Violet 421 | 323219 | 43A3 |
| Biolegend | Anti-CD44 PE | 103023 | IM7 |
| Biolegend | Anti-CD73 PE/Dazzle 594 | 344019 | AD2 |
| Biolegend | Anti-CD106 APC | 305809 | STA |
| Biolegend | Anti-CD36 PE-Cy7 | 336221 | 5-271 |
| Biolegend | Anti-CD45 FITC | 304005 | HI30 |
| Biolegend | Anti-CD34 FITC | 343503 | 581 |
| Biolegend | Anti-CD11b FITC | 301329 | ICRF44 |
| Biolegend | Anti-CD19 FITC | 302205 | HIB19 |
| Biolegend | Anti-HLA DR FITC | 307603 | L243 |

### Trilineage Differentiation Assays

For adipogenic induction, we cultured P3 BMSCs on 0.1% gelatin-coated plates at 37°C/5%CO_2_/5%O_2_ in alpha-MEM (Gibco, ThermoFisher, MA) supplemented with 15% fetal-bovine-serum (FBS) (Gibco, ThermoFisher, MA), 1% non-essential amino acids (Gibco, ThermoFisher, MA), and 1% P/S (Gibco, ThermoFisher, MA), 0.5uM dexamethasone, 0.5mM IBMX, 50uM Indomethacin, and 1.5 mM insulin. We changed media every 2 days for 21 days. We assayed for adipogenic differentiation by with Oil-Red-O that preferentially dissolves in lipid droplets. For osteogenic induction, we cultured P3 BMSCs at 37°C/5%CO_2_/5%O_2_ in alpha-MEM (Gibco, ThermoFisher, MA) supplemented with 15% fetal-bovine-serum (FBS) (Gibco, ThermoFisher, MA), 1% non-essential amino acids (Gibco, ThermoFisher, MA), and 1% P/S (Gibco, ThermoFisher, MA), 100nM dexamethasone, 50μM ascorbate phosphate, 10mM β-glycerophosphate and 100ng/mL BMP-2 (Peprotech, NJ), that we changed every 2 days for 14 days. We assayed osteogenic differentiation for calcium deposits using alizarin red staining. We photographed stained cells using a Zeiss Observer microscope.

For chondrogenic induction, we pelleted 2.5×10^5^ BMSCs in 15mL tubes at 200 *xg* for 5 minutes. We then cultured the cell pellets at 37°C/5%CO_2_/5%O_2_ in alpha-MEM containing 15% FBS, 1% non-essential amino acids (Gibco, ThermoFisher, MA), 1% P/S, 100nM dexamethasone, 50μg/mL ascorbic acid 2-P, 40μg/mL L-Proline, 1% ITS/supplement B-D, 1mM sodium pyruvate, 10ng/mL TGFβ3 (Peprotech, NJ) and 100ng/mL BMP2 (Peprotech, NJ) for 21 days. We fixed pellets in 4% paraformaldehyde (PFA), embedded in OCT, sectioned and stained for cartilage using Safranin-O with Hematoxylin R and Fast Green for counterstaining.

### Exosome prep

We isolated exosome preps from conditioned media, of the plastic adherent human BMSCs. We adapted a published protocol(Théry et al., 2006) with modifications to optimize yield and quality of exosomes from our cell type and culture conditions. Following a series of differential centrifugations to remove dead cells, cell debris and medium/large vesicles, we filtered the supernatant through a 200nm PES filter (**Supplemental Figure 1**). We pelleted exosome preps at 100,000 *xg* for 90 minutes in a Sorvall Wx ultracentrifuge using ThermoFisher F37L fixed angle rotors. We consolidated exosome prep pellets, resuspended in PBS and re-centrifuged at 100,000 *xg* for 90 minutes for a final pellet using a Beckman Coulter 70 Ti rotor. Following resuspension in 1x PBS, we stored exosome prep aliquots in -80°C until use, without any freeze/thaw cycles.

### Nanoparticle Tracking Analysis

To measure the hydrodynamic diameter and concentration of BMSC exosome preps, we used nanoparticle tracking analysis. Using a Zetaview instrument and software (Particle Metrix, Germany), we injected 1mL exosome prep sample into the flow cell and captured videos at 30 frames/second, at 11 different imaging positions. For statistic validity, we only used samples and readings that had particles in 10 out of 11 positions (manufacturer’s guidelines are that ≥ 7 positions necessary for validity). The built-in algorithm uses the light scattering, as well as drift or Brownian motion of the particles in a Stokes-Einstein equation to calculate particle size. We ensured capture and analysis of minimum 1000 traces of particles per run for characterization and quantification. Our post-processing parameters were max area 1000, min area 10, min brightness 20, tracelength 30.

### Exosome prep tetraspanin composition analysis

We submitted exosome preps to NanoView Biosciences (now part of Unchained Labs, CA) for analysis of tetraspanin proteins CD63, CD81 and CD9. Per manufacturer’s notes, capture and fluorescence-conjugated detection antibodies are of the same clones for each antibody, with CD63, CD81 and CD9 fluorescently labeled with CF-647, CF-555 and CF-488, respectively. The results incorporate ≥3 technical replicates. We used the interferometric measurements and fluorescent tetraspanin counts directly exported from the ExoView Analyzer software and, if necessary, visualized further using GraphPad Prism 9 (Dotmatics, MA).

### Exosome prep labeling with fluorescence

We labeled exosome preps with ExoGlow-Vivo EV Labeling Kit (System Biosciences, CA) per manufacturer’s instructions. Briefly, we incubated exosome preps with ExoGlow-Vivo NIR dye for 1 hour at room temperature. We removed unconjugated ExoGlow-Vivo dye by washes in 1xPBS w/Ca^2+^Mg^2+^ and ultracentrifugation at 100,000*xg*. We resuspended exosome prep pellets in 1xPBS w/Ca^2+^Mg^2+^ and proceeded to encapsulation with Q. Alternatively, we labeled using ExoGlow-RNA EV Labeling Kit (System Biosciences, CA) for GFP fluorescence.

### Q protein biosynthesis

We expressed and purified Q as published(Hume et al., 2014). Briefly, we transformed ampicillin-resistant pQE30/Q plasmids into chemically competent M15MA *Escherichia coli* cells. We grew the transformed cells in supplemented minimal M9 media and induced by adding isopropyl ß-D-1-thiogalactopyranoside (IPTG). We pelleted Q expressing-cells, resuspended and sonicated in 40 mL of buffer A (50 mM Tris-HCl, 500 mM NaCl, pH 8). Based on the hexahistidine tag on the N-terminus of Q, we purified Q using syringe-pump driven immobilized metal affinity chromatography with a cobalt-charged Q sepharose fast-flow 5 mL column. Following elution using a gradient of buffer B (buffer A, 500 mM imidazole), we collected and dialyzed pure fractions against buffer A (6×5L for 4 hours each) to remove imidazole.

### Hydrogel Preparation

We concentrated Q to 2 mM via centrifugal filters with a 3 kD MWCO at 3000 rpm for 30 minutes intervals. After confirming concentration in BCA assays, we incubated 250 µL aliquots of Q at 4°C in a 2 mL glass vial and visually inspected at different timepoints to gauge gelation; tube inversion tests confirmed gelation.

### Rheology

We assessed the rheological properties of Q with Exo on a stress-controlled rheometer, equipped with 8 mm parallel plates set to a 0.2 mm geometry gap and a temperature of 4°C. Strain sweeps in a range of 0.1 to 1000% strain at a frequency of 1 Hz determined the linear viscoelastic region of Q. We carried out frequency sweeps from 0.1 to 1.

### Transmission and scanning electron microscopy of hydrogels

We diluted gel samples from 2 mM to 50 uM and spotted 3 µL immediately on Formvar/carbon-coated copper grids. After blotting the samples with filter paper, we washed with 5 µL dH2O, blotted again, and then treated with 3 µL 1% uranyl acetate for 1 min to negatively stain the sample followed by blotting. We acquired TEM images at an accelerating voltage of 120 kV using Philips CM12 electron microscope. We spotted 5 µL gel samples onto a silica wafer and allowed to dry in a vacuum desiccator for 3 hours. We then iridium-coated the samples to 3 nm using a high-resolution sputter coater, prior to SEM image acquisition using MERLIN field emission scanning electron microscope (Carl Zeiss AG).

### Mouse diabetic wound model and data analysis

C57BL/6J and BKS.Cg4Dock7^m+/+^LepR^db/J^ strains (Jackson Laboratories) are housed in a clean, daylight facility and fed mouse chow ad libitum. The IACUC at New York University School of Medicine approved all protocols. We used 12-16 weeks old LepR^db/db^ type 2 diabetic mice, with blood glucose ≥350 mg/dL for all studies to ensure the mice manifest all severe complication of diabetes. Under inhalational anesthesia, we clipped the dorsal hair and used Nair to clear all hair shafts. Then we used a 10 mm biopsy punch on mouse dorsal skin to create a full-thickness (excisional) wound. This wound technique removes all layers of the skin – cornified layers, epidermis, dermis, adipose tissue, and panniculus carnosus – without compromising the underlying fascia. To simulate human-like healing by secondary intention, a silicone stent placed around the excisional circle and sutured into place with interrupted sutures prevents premature contraction from the panniculus carnosus muscle layer in mouse skin, that is absent in humans (Galiano et al., 2004; Rabbani et al., 2018; Short et al., 2023). Non-stented mouse wounds close primarily by contraction of the panniculus carnosus and not via granulation tissue, unlike human skin tissue repair. We placed a clear, plastic blister pack to maintain moisture during healing, ensuring no contact of the plastic cover with the wound or hydrogels. We provided analgesics for 72 hours post-op. We applied one dose of hydrogels at POD1. We monitored wounds and scabs with photometric measurements and calculated open wound area as a percentage of the original POD0 wound, identical in size to the 10mm diameter silicone stent. Pathological time to closure is time in excess of physiologic times in WT mice. We calculated wound burden by integrating the area under the curve, open wound area against time, using the trapezoid rule (Margolis et al., 2002).

### In vivo Exo-Q hydrogel imaging

To capture epi-fluorescence from mice, we used an In Vivo Imaging System (IVIS) Perkin Elmer IVIS Lumina III for NIR ex/em 740/845nm (Waltham, MA) or Bruker In Vivo Xtreme for GFP 480/535nm (Carteret, NJ). With mice under inhalational anesthesia, we captured any fluorescence from the fluorescently labeled EV preps with the ultrasensitive CCD camera in the IVIS. We used Living Image (Perkin Elmer, Waltham, MA) to analyze images.

### Cardiac injection of lectin

We injected Dylight-labeled tomato lectin (592/617nm, Vector Labs, CA) through cardiac injection into anesthetized mice. After 5 minutes, based on Robertson et al (Robertson et al., 2015), we flushed the vascular system by cardiac perfusion with 10mL 1X PBS. We monitored the outflow from the ligated vena cava, as well as liver color and tail curvature to ensure appropriate perfusion. Then we perfused with 4% PFA, 5mL/min. Then we collected the wounds and other organs, fixed further by submersion in 4% PFA for 48 hours at 4°C.

### Wound tissue collection

We collected wound tissues, bisected in cranial-caudal orientation. Following overnight fixation in 4% PFA pH 7.0 at 4°C, skin tissues underwent dehydration into paraffin blocks for H&E and Opal immunofluorescent staining by the Experimental Pathology Laboratory. We used 5µm tissue sections for analysis. For wound tissues with fluorescent labeled exosomes or lectin, we cryopreserved samples in 15% and then 30% sucrose, then prepared cryoblocks in OCT for 10um sections. We used DAPI to counterstain the tissue sections.

### Statistical analyses

All data is represented as mean ± standard deviation, n≥3 biological replicates. We used one-way ANOVAs when comparing 3 or more sets of data and either Tukey or Dunnet post-hoc corrections, when comparing samples against one control or every sample to another, respectively. Based on prior work, we used GPower (Düsseldorf, Germany) to determine that we needed minimum 4 diabetic wounds per condition for an effect size of 5.

### RNASeq and analysis

#### Exosomes

We submitted exosome preps that were stored at -80°C, on dry ice to System Biosciences (Palo Alto, CA) for their Exosome RNA next-generation sequencing (NGS) service, Exo-NGS. Per the provider, small RNA libraries are constructed with the CleanTag Small RNA Library Preparation Kit (TriLink, Cat# L-3206) according to the manufacturer’s protocol. The purified library is quantified with High Sensitivity DNA Reagents (Agilent Technologies, Part # G2933-85004) and High Sensitivity DNA Chips (Agilent Technologies, Part # 5067-4626). The libraries are pooled and to ensure optimal size the 140bp to 300bp region is size selected on an 8% TBE gel (Invitrogen by Life Technologies, Ref# EC6215). The size selected library is quantified with High Sensitivity DNA 1000 Screen Tape (Agilent Technologies, Part # 5067-5584), High Sensitivity D1000 reagents (Agilent Technologies, Part# 5067-5585), and the TailorMix HT1 qPCR assay (SeqMatic, Cat# TM-505), followed by a NextSeq High Output single-end sequencing run using the NextSeq 500/550 High Output v2 kit, following manufacturer’s instructions.

#### Cells

The NYU Genome Technology Center at the NYU School of Medicine generated total RNA-Seq libraries for human BMCSs. Samples underwent rRNA depletion and library construction using Nugen Trio RNA Seq (Tecan Genomics, #0606-96), followed by sequencing using the Illumina NovaSeq 6000 and SP-100 flow cell sequencing kit (Illumina, 20027464). The settings for the sequencing run were configured to 50PE with a target read depth of 0.5X. Each library was assigned to a separate flow cell lane and sequenced accordingly. The reads were then analyzed either individually or collectively from each lane.

#### Sequence processing

We then determined adapter trimming and quality control parameters using FASTQC analysis of cell and exosome preparations. We performed trimming for the pair-end RNA seq data prior to alignment, retaining reads with high-quality scores and removing adapters. We trimmed single-end reads from the exosome preps via TrimGalor based on the small non-coding RNA expected size (J. Li et al., 2020). After trimming, we align the reads to the hg38 human genome using STAR and utilize optimization of the alignment for small non-coding RNA parameters with uniquely mapped reads over 70%. Several custom parameters are added to the ENCODE small-RNA-seq pipeline to minimize multi-mapping and unmapped reads (Dunham et al., 2012). We set –outFilterMismatchNoverLmax at 0.05 to retain alignment quality across all the reads, –winAnchorMultimapNmax at 1000 to improve the mappability, and –outFilterMatchNmin at 16 to retain as many small non-coding RNA in our data. We used a customized GTF file with DASHR, RNAcentral, miRbase, and Ensembl to annotate samples. The following BAM files then went through the MGcount pipeline to account for multialignment, multimapping, and annotation sources (Hita et al., 2022). The output results are a counts matrix, feature metadata, and a multimap graph — the DEseq analysis of the count matrix with a design model set to cells or exosomes. We performed subsequent variance stabilization transformation and scaled by column to generate the PCA plot. To generate the heatmap, we log transform all samples and generate the top 30 significantly differentiated genes based on |log2FC| > 0.5 and adj pval <0.1 and scale by row.

### Quantitative RT-PCR

We homogenized whole wound tissues in Trizol (ThermoFisher) using a bead ruptor (Fisherbrand™ Bead Mill 24), then transferred to fresh tubes for chloroform-based separation. We transferred the aqueous phase to fresh tubes and precipitated RNA with iso-propyl alcohol (ThermoFisher), followed by purification using spin columns of RNeasy Mini kits (Qiagen). We used 500ng total RNA for reverse transcription using the High-Capacity cDNA synthesis kit (ThermoFisher) and quantified in real-time with SYBRgreen detector in a QuantStudio7-Flex (ThermoFisher). We used the delta-delta-CT method to calculate relative expressions.

### Xenograft wound model

We performed xenografts adapted from published methods (Karim et al., 2020). Briefly, we obtained de-identified abdominal human skin from patients undergoing plastic reconstructive procedures, such as panniculectomies, using an Institutional Review Board exempt protocol in accordance with the regulations of New York University Langone Health. We transported the tissue on ice to the mouse surgery area, removed subcutaneous fat with scissors, rinsed in normal saline and maintained 2cm x 1cm grafts in DMEM in 100mm petri dishes on ice. We removed 2cm x 1cm dorsal skin from 8-10 weeks old Rag1^-/-^ mice (B6.129S7-*Rag1^tm1Mom^*/J, Jackson Laboratories, ME), placed the human skin graft and sutured at the four cardinal positions using 6-0 vicryl sutures, ensuring the skin are everted. We placed one suture in each side of the graft to approximate gaps and dressed the graft with petroleum jelly, covered with sterile gauze and secured to the body of the mouse with elastic, adhesive wraps. We provided analgesics for 72 hours post-op and changed bandages every week for 4 weeks. We used size 15 blades to create 5mm x 5mm wounds in the xenografts.

