## Supplemental Materials for "Functional high capacity exosome-encapsulating bioinspired hydrogel promotes microvascular bed expansion in diabetic mice"

Supplemental Figures 1-9

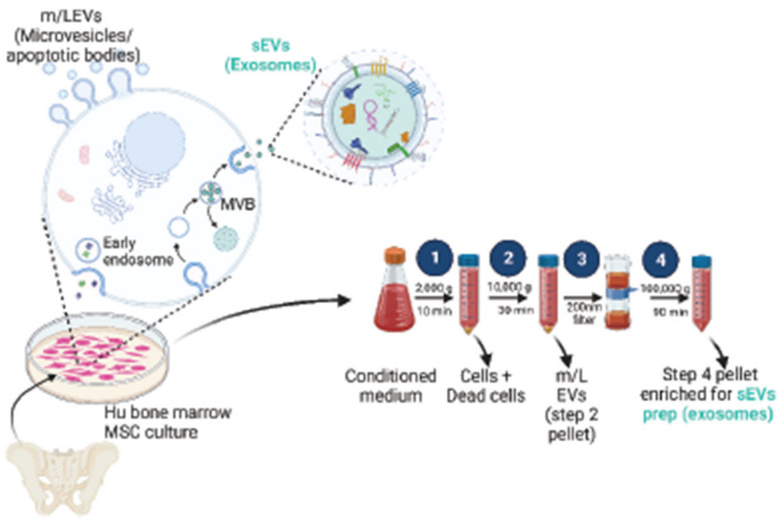


**Supplementary Figure 1** (related to Figure 1 and Methods): Exosome prep isolation steps from human BMSC conditioned media. Schematic created with BioRender.com.

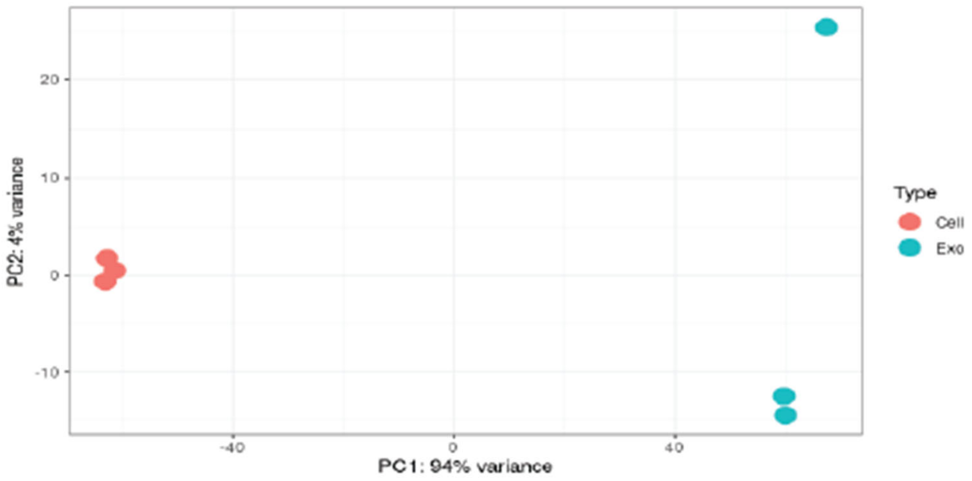


**Supplementary Figure 2** (related to Figure 2): Regularized log transformation PCA plot of Cells vs Exosomes

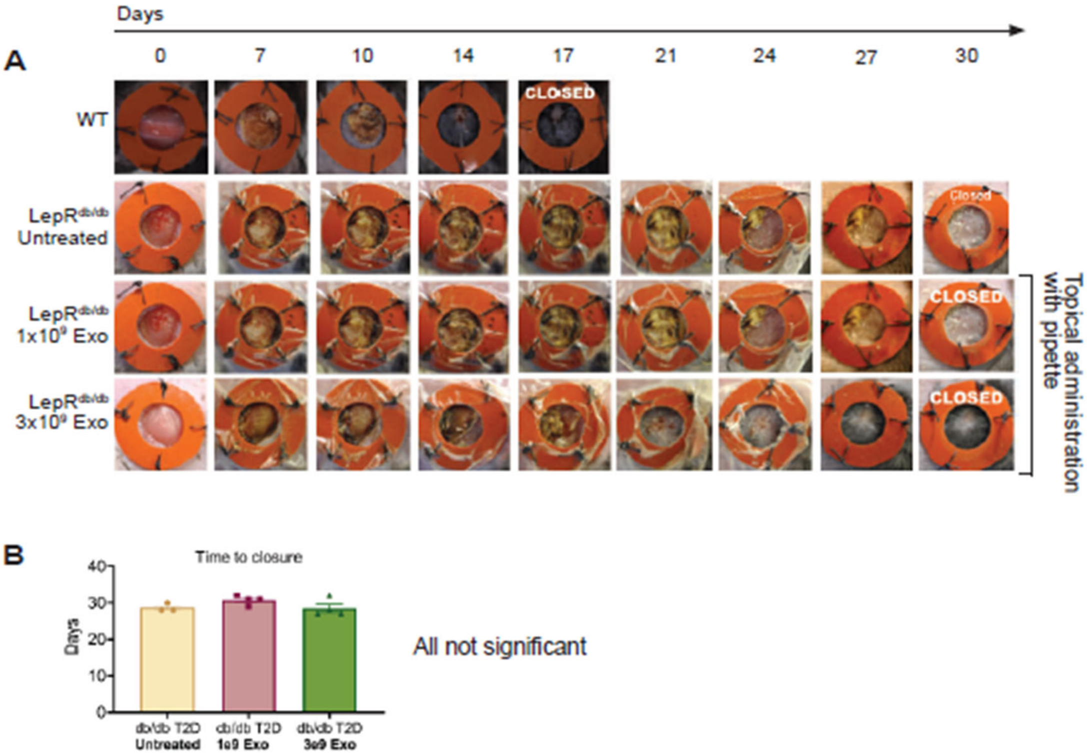


**Supplementary Figure 3** (related to Figure 3): Topical application of exosome suspension, drop by drop, with pipettes does not affect diabetic wound closure times. a) Wound photographs, monitored from excision until closure, with exosome administration by pipetting onto wounds at POD1. b) Mean time to closure. Data is represented as mean ± standard deviation.

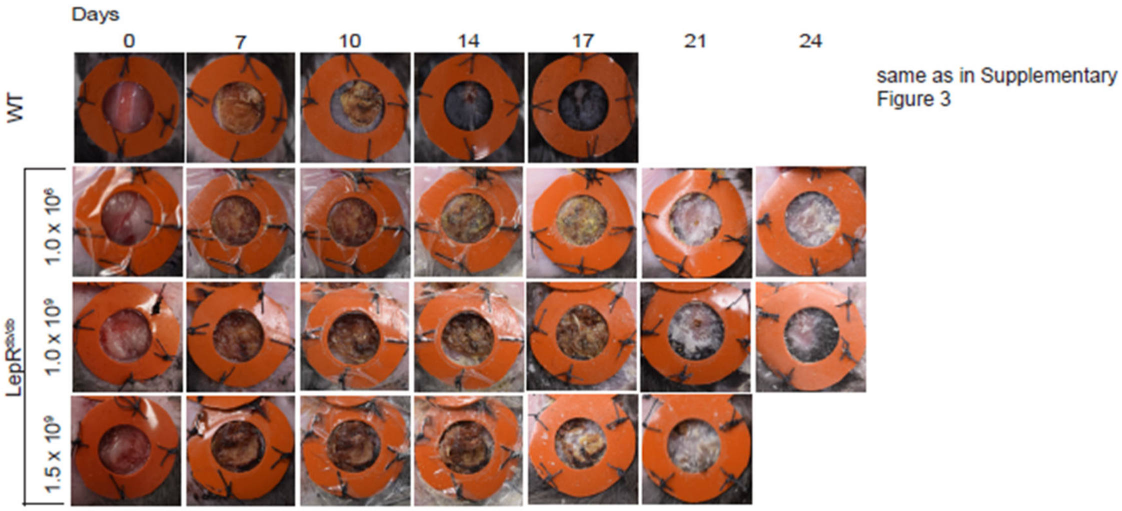


Supplementary Figure 4 (related to Figure 4): Time to closure assessment with varying dose of exosomes. Wounds photographed, monitored after exosome administration in hydrogels at POD1.

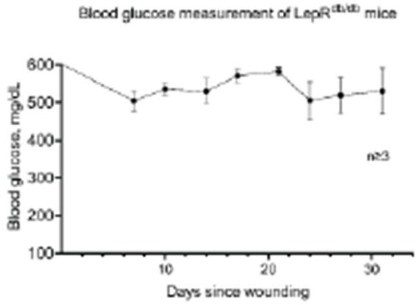


**Supplementary Figure 5** (related to Figure 4): Hyperglycemia is not affected and is maintained in LepR$^{db/db}$ type 2 diabetic mice throughout wound closure time course, regardless of treatment modality.

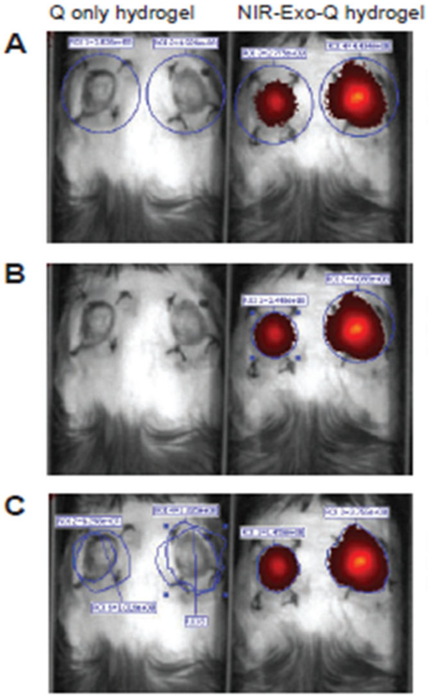


A: Fixed diameter ROI around each wound - measures fluoresence intensity within the circles of identical area.

B: Maximum diameter ROI around each wound - measures largest circular area that captures fluoresence signal area.

C: Planar area ROI around fluorescent signal - measures hand-drawn area fitted around fluoresence signal.

**Supplementary Figure 6** (related to Figure 6): Quantification methods for epi-fluorescence-based in vivo detection of NIR-Exo-Q hydrogel. Region of interest (ROI) allows identification of different wounds. Copy of identical shaped ROI is for calculation of fluorescence intensity of control-treated wound.

Supplementary Figure 7 (related to Figure 7): Wound tissue sections from POD5. H&E and multiplex immunostains as indicated. ). Dashed white lines, epidermal-dermal boundary. Solid white line, granulation tissue-dermis boundary. Ɛ, epidermis. Δ, wound-adjacent dermis. §, granulation tissue. Scale bar, 200µm.

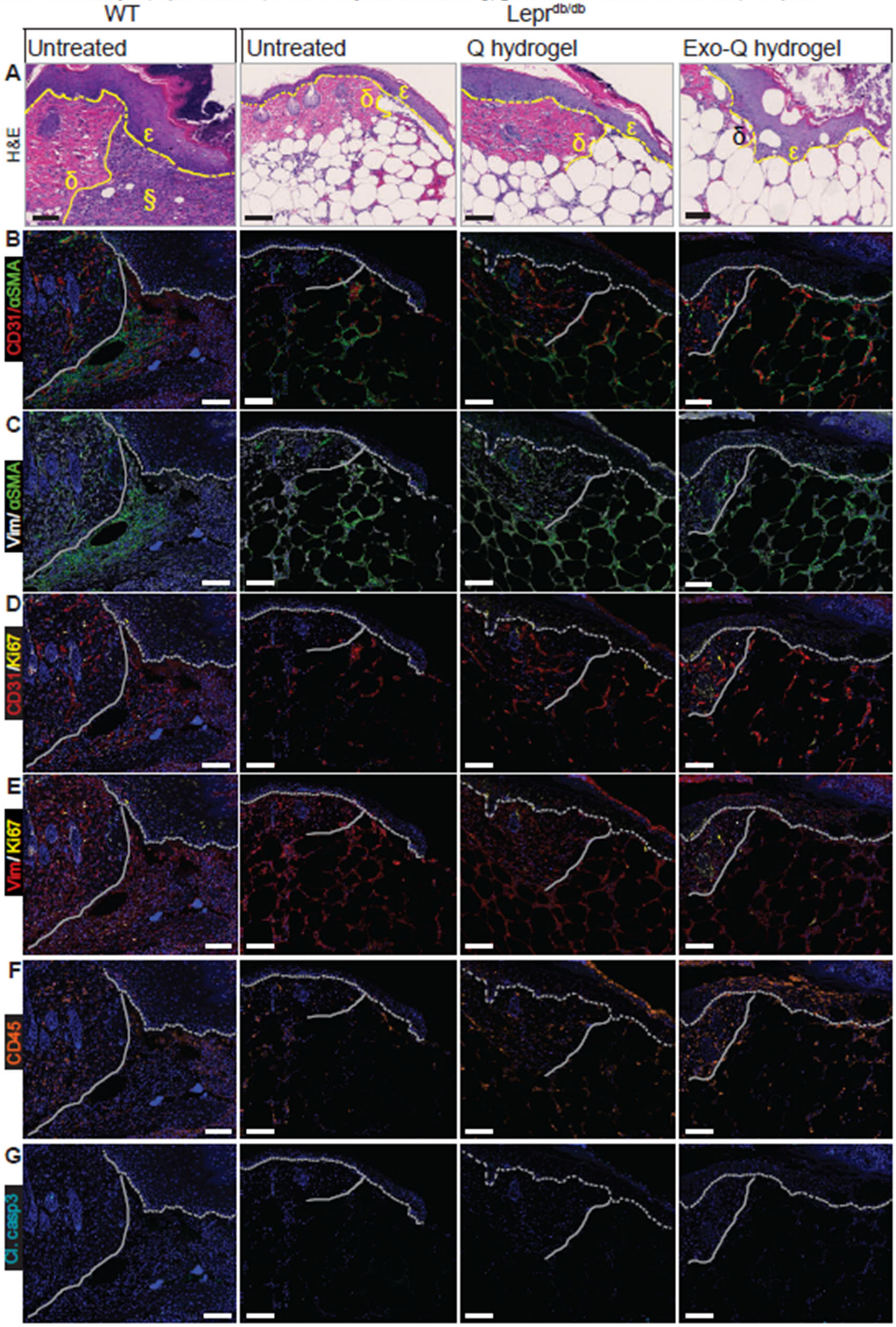

Week 6 Week 33

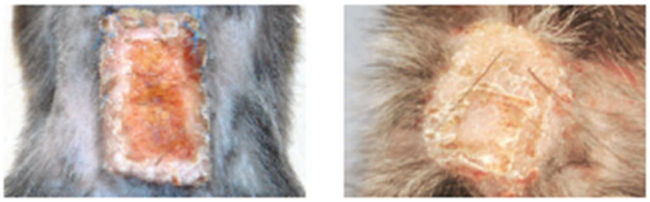

**Supplementary Figure 8** (related to Figure 8): Human skin xenografts at indicated timepoints post-grafting on Rag1$^{-/-}$ mice. Human hair regrowth, photographed at Week 33 indicates xeno-graft vascular anastomoses with mouse vasculature and graft survival.

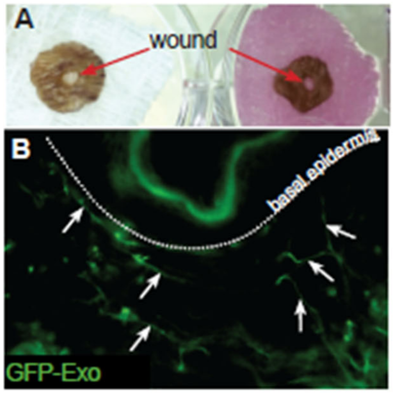


**Supplementary Figure** 9 (related to Figure 8): Human skin explants, with topical administration of Exo-Q hydrogel, containing GFP-labeled Exo. Transduction of immortalized adult adipose multipotent stromal cells (ADMSC) with a CD81-GFP-coding lentivirus allowed generation of CD81-GFP-labeled ADMSC Exo. A) Human skin explants with "wounds". The skin explant tissue is placed on sterile gauze (left) and maintained on culture media (right) for the duration of the experiment. B) GFP signal detection in fixed tissue sections of human explants with wounds. Dashed line indicates epidermal-dermal border. Photographs are of wound-adjacent areas. The GFP signal is consistent with appearance of microvasculature of the human skin explant.
